# Inferring local movement of pathogen vectors among hosts

**DOI:** 10.1101/046540

**Authors:** Z Fu, B Epstein, J. L. Kelley, Q. Zheng, A. O. Bergland, C. I. Castillo Carrillo, A. S. Jensen, W. E. Snyder

## Abstract

Herbivores often move among spatially interspersed host plants, tracking high-quality resources through space and time. This dispersal is of particular interest for vectors of plant pathogens. Existing molecular tools to track such movement have yielded important insights, but often provide insufficient genetic resolution to infer spread at finer spatiotemporal scales. Here, we explore the use of <u>Nex</u>tera-<u>t</u>agmented <u>r</u>eductively-<u>a</u>mplified <u>D</u>NA (NextRAD) sequencing to infer movement of a highly-mobile winged insect, the potato psyllid (*Bactericera cockerelli*), among host plants. The psyllid vectors the pathogen that causes zebra chip disease in potato (*Solanum tuberosum*), but understanding and managing the spread of this pathogen is limited by uncertainty about the insect’s host plant(s) outside of the growing season. We identified 8,443 polymorphic loci among psyllids separated spatiotemporally on potato or in patches of bittersweet nightshade (*S*. *dulcumara*), a weedy plant proposed to be the source of potato-colonizing psyllids. A subset of the psyllids on potato exhibited close genetic similarity to insects on nightshade, consistent with regular movement between these two host plants. However, a second subset of potato-collected psyllids was genetically distinct from those collected on bittersweet nightshade; this suggests that a currently unrecognized host-plant species could be contributing to psyllid populations in potato. Oftentimes, dispersal of vectors of plant or animal pathogens must be tracked at a relatively fine scale in order to understand, predict, and manage disease spread. We demonstrate that emerging sequencing technologies that detect SNPs across a vector’s entire genome can be used to infer such localized movement.

## Introduction

Herbivores often move among host plant species, driven by their need to evade and detoxify plant defenses, balance nutritional requirements that cannot be met by single plants, and/or track spatiotemporal variation in plants’ resource quality (Fryxell & Sinclair 1988; Loxdale & Lushai 1999). At the broadest scale, herbivores may traverse thousands of kilometers, tracking host-plant availability across seasons (e.g., Despland *et al*. 2004; Bischof *et al*. 2012; Chasen *et al*. 2014; Shariatinajafabadi *et al*. 2014) or due to varying rainfall and wind patterns (e.g., Rose 1979; Zhu *et al*. 2006). At a finer scale, herbivores often move among host-plant species within a habitat while tracking host-plant phenology, as different host-plant species go through seasonal changes in nutritional value and/or ability to physically or chemically defend themselves (Peterson 1997; Wilmshurst *et al*. 1999; Monteith *et al*. 2011). When herbivores act as vectors of plant pathogens, these movements can have particularly dramatic effects on host plants; herbivores can initiate pathogen outbreaks even when herbivore densities are too low to inflict appreciable direct damage (Nault 1997; Redak *et al*. 2004; Weintraub and Beanland 2006). Oftentimes, a detailed understanding of movement of vectors among host plant species, or within stands of the same species, is critical for predicting patterns of disease spread (e.g., Power 1987; McElhany *et al*. 1995; Weintraub & Beanland, 2006).

When herbivores are relatively large, or the distances covered are relatively small, physically marking and tracking individual herbivores can be an effective way to unravel patterns of host-plant switching (Bach 1980; Belovsky 1981; Illius *et al*. 1992; Hagler & Jackson 2001). However, when this is impossible or impractical, patterns of interrelatedness among herbivores can be used to infer likely movement patterns. Molecular techniques, including protein and microsatellite DNA markers, were among the first genetic tools used to infer gene flow and thus herbivore dispersal (Loxdale & Lushai 1998; Behura 2006; Wink 2006). However, developing a sufficiently large set of markers to delineate localized movement can be time consuming and expensive, or even impossible when there are few microsatellites in the genome (Zhang 2004; Meglécz *et al*. 2007; Coates *et al*. 2009). Most recently, restriction-site associated DNA (RAD) markers have been used to overcome these limitations by allowing relatively quick identification of single nucleotide polymorphisms (SNPs) across focal organisms’ entire genomes. RAD-based approaches have proven powerful in tracking genetic differentiation across landscapes (e.g., Barley *et al*. 2015; Szulkin *et al*. 2016), but the relatively high DNA-volume inputs required by these techniques has thus far limited their use to larger-bodied organisms. Because of their small body sizes, many herbivorous insects that feed heavily on plants (and/or vector key plant pathogens) have thus far been outside the reach of these approaches.

Here, we explore the use of Nextera-tagmented reductively-amplified DNA (“NextRAD”) sequencing to infer movement among host plant species by a winged, small-bodied insect, the potato psyllid (*Bactericera cockerelli*). The psyllid is the vector of the bacterium (*Candidatus* Liberibacter solanacearum) that causes zebra chip disease in cultivated potato (*Solanum tuberosum*) (Liefting et al. 2009), whose spread has endangered potato production in several parts of the world (e.g., Greenway 2014). In the northwestern U.S., it has been proposed that potato psyllids transmit the zebra chip pathogen as the insects migrate from the perennial solanaceous weed bittersweet nightshade, *Solanum dulcumara*, to annually-cultivated *S*. *tuberosum* fields each year (Horton *et al*. 2015a). However, movement of potato psyllids from bittersweet nightshade to potato has never been directly demonstrated, hindering any ability to understand, predict, or manage zebra chip outbreaks (Horton *et al*. 2015a). Indeed, a wide variety of plant species other than bittersweet nightshade has been proposed to be the true source of psyllids (and perhaps also the zebra chip pathogen) that colonize potato fields (Horton *et al*. 2015b). Existing molecular tools for dividing psyllids into geographically-separated genetic groups, based on sequence variation within the Cytochrome *c* Oxidase I (COI) gene, are too limited to reveal genetic subpopulations at a fine enough scale to identify movement among host plants (Horton et al. 2015a). NextRAD sequencing overcomes these limitations by fragmenting and ligating adaptor sequences to genomic DNA via engineered transposomes (Nextera DNA Library Prep Reference Guide), detailing genetic differentiation across the insect’s genome. Critically, NextRAD requires less than 50 ng of DNA (Russello *et al*. 2015; Nextera DNA Library Prep Reference Guide) making it possible to provide sequence data from organisms far smaller than was possible with RAD sequencing approaches. Using this technique, we assessed whether bittersweet nightshade could be the sole source of potato psyllids colonizing potato fields, or whether instead other non-crop host species might need to be identified. This approach may have wide applicability, helping to understand spread of the many plant and animal pathogens vectored by small, highly-mobile insects.

## Materials and Methods

Our project included regional sampling of spatially-dispersed herbivore populations on two host plant species, followed by sequencing the insects to infer population interrelatedness. First, over two years, we collected potato psyllids from bittersweet nightshade patches located throughout much of the potato-growing region of east-central Washington State (USA); in one of these years, we also collected psyllids from production potato fields across this same region (Fig. 1). Additionally, we collected psyllids from a nightshade patch located in southern Idaho (Fig. 1) to serve as a geographically-distinct outgroup. A subsample of the psyllids collected from nightshade patches (up to 10 psyllids per sampling date), and all psyllids collected from potato fields, were then sequenced using the NextRAD approach; this allowed us to identify variant sites throughout the psyllid genome. We then used multiple population-genetic approaches to determine interrelatedness among psyllids collected from the two host plants, to infer whether they are composed of a single interbreeding population or instead include members of genetically-distinct sub-populations. This allowed us to infer whether the bittersweet nightshade patches that we sampled could represent the sole source of our potato-collected psyllids. Each of these project sub-components are detailed below.

**Fig. 1.**
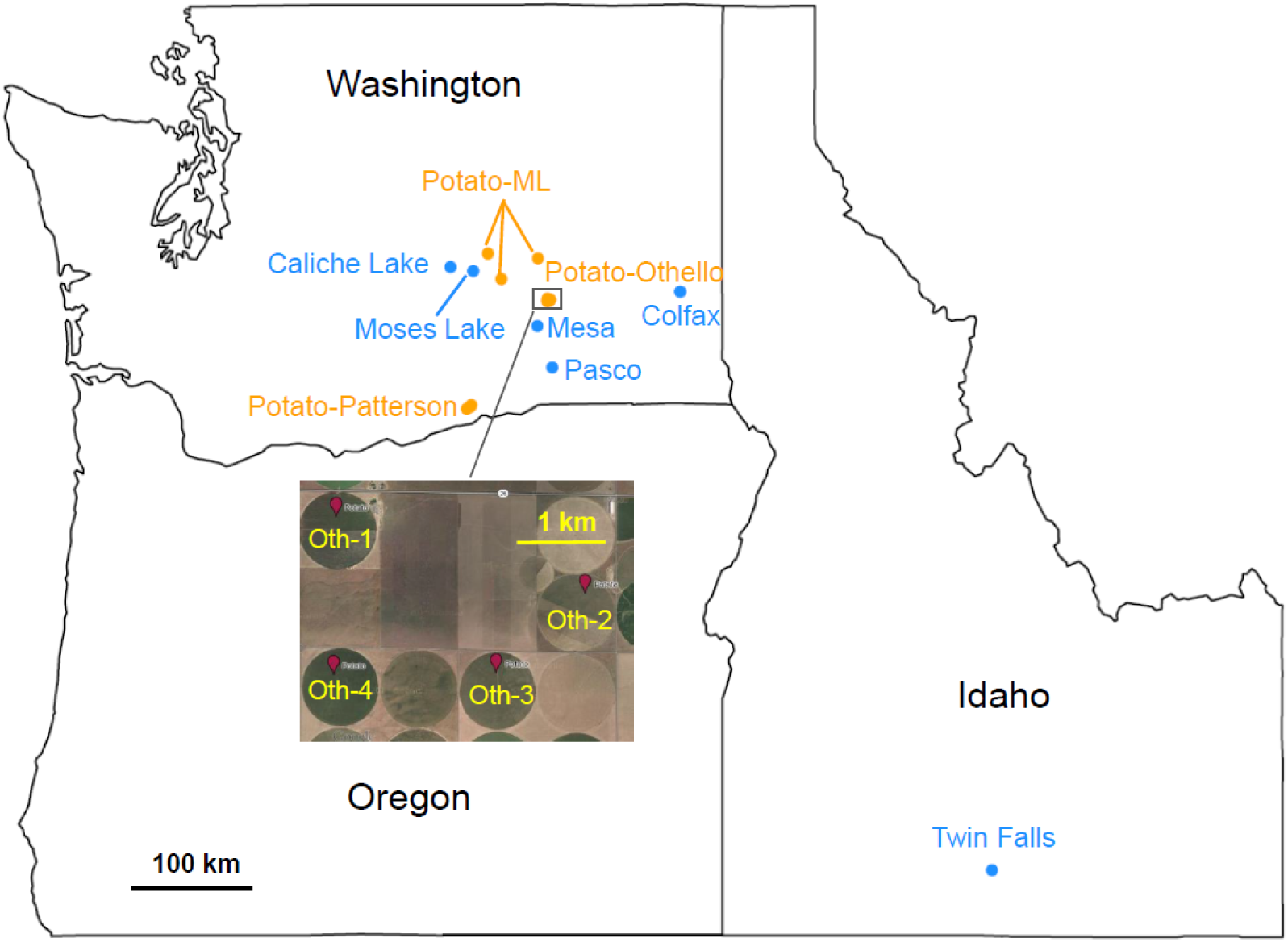
Potato psyllid collection sites across the states of Washington and Idaho, USA (see Table 1 and Table S1, Supporting Information, for details on collection sites and dates). Psyllids were collected from bittersweet nightshade patches (blue circles) or potato fields (yellow circles).

### Potato psyllid sampling and sequencing

Potato psyllids were collected from six bittersweet nightshade patches and ten potato fields in the U. S. states of Washington and Idaho (Table 1, Fig. 1), over two growing seasons, using a suction sampling device (see Koss *et al*. 2005; Crowder *et al*. 2010). Sites were chosen to cover the majority of the potato-growing region in east-central Washington, with the Idaho site serving as a geographically-distant outgroup, and were sampled periodically over the 2012 and 2013 growing seasons (Fig. 1, Table S1, Supporting Information). Psyllids were placed on dry ice immediately following collection, and were stored in 95 % ethanol upon arrival in the laboratory. For each sampling date, on each host plant and at each location considered here, up to ten intact adult psyllids were randomly selected for DNA extraction and sequencing (Table 1, Table S1, Supporting Information). In total, we processed 285 psyllids for NextRAD sequencing.

**Table 1.**
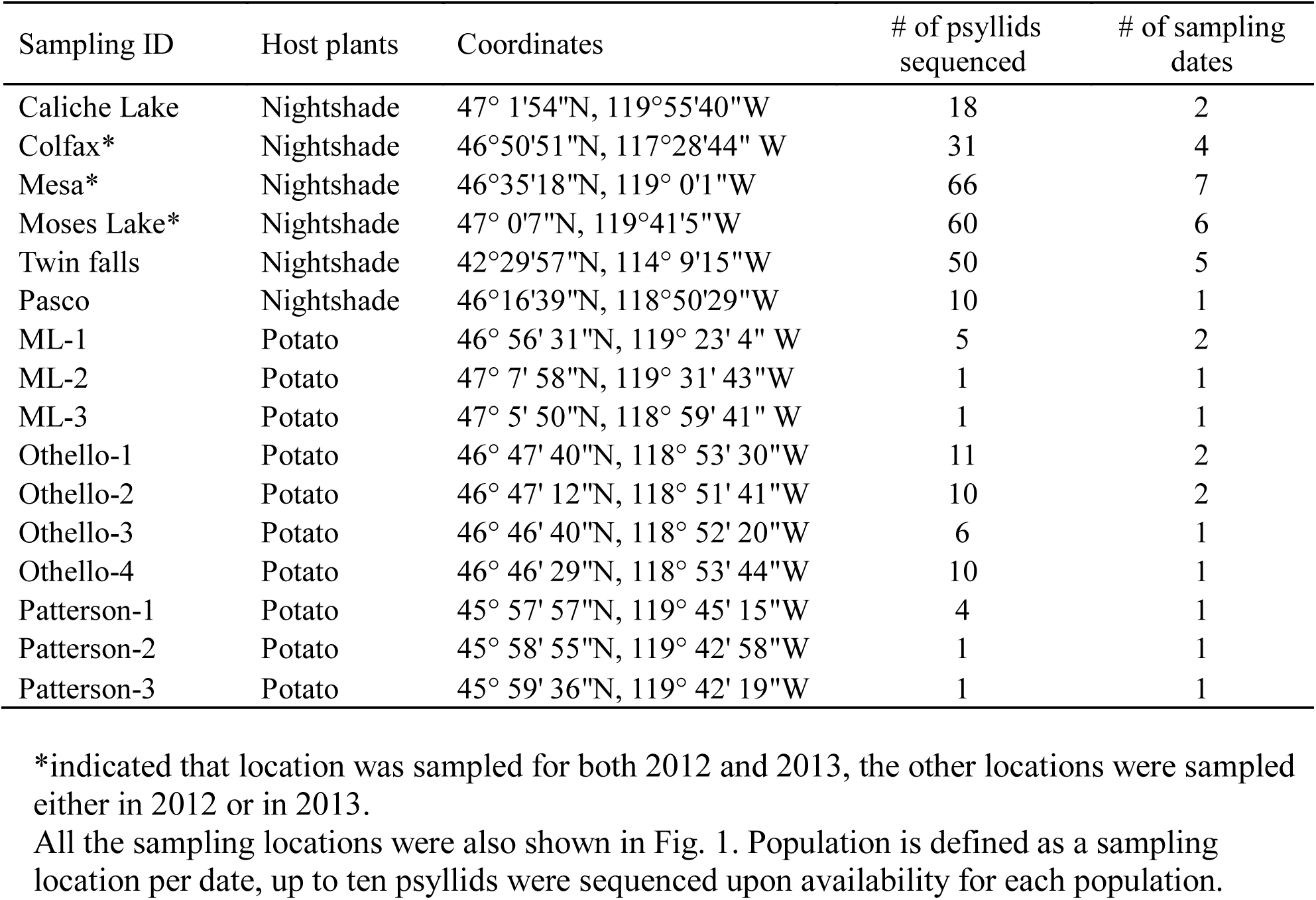
Sampling locations and sampling dates of potato psyllids *Bactericera cockerelli* in the states of Washington and Idaho, USA.

| Sampling ID | Host plants | Coordinates | # of psyllids sequenced | # of sampling dates |
| --- | --- | --- | --- | --- |
| Caliche Lake | Nightshade | 47° 1'54"N, 119°55'40"W | 18 | 2 |
| Colfax* | Nightshade | 46°50'51"N, 117°28'44" W | 31 | 4 |
| Mesa* | Nightshade | 46°35'18"N, 119° 0'1"W | 66 | 7 |
| Moses Lake* | Nightshade | 47° 0'7"N, 119°41'5"W | 60 | 6 |
| Twin falls | Nightshade | 42°29'57"N, 114° 9'15"W | 50 | 5 |
| Pasco | Nightshade | 46°16'39"N, 118°50'29"W | 10 | 1 |
| ML-1 | Potato | 46° 56' 31"N, 119° 23' 4" W | 5 | 2 |
| ML-2 | Potato | 47° 7' 58"N, 119° 31' 43"W | 1 | 1 |
| ML-3 | Potato | 47° 5' 50"N, 118° 59' 41" W | 1 | 1 |
| Othello-1 | Potato | 46° 47' 40"N, 118° 53' 30"W | 11 | 2 |
| Othello-2 | Potato | 46° 47' 12"N, 118° 51' 41"W | 10 | 2 |
| Othello-3 | Potato | 46° 46' 40"N, 118° 52' 20"W | 6 | 1 |
| Othello-4 | Potato | 46° 46' 29"N, 118° 53' 44"W | 10 | 1 |
| Patterson-1 | Potato | 45° 57' 57"N, 119° 45' 15"W | 4 | 1 |
| Patterson-2 | Potato | 45° 58' 55"N, 119° 42' 58"W | 1 | 1 |
| Patterson-3 | Potato | 45° 59' 36"N, 119° 42' 19"W | 1 | 1 |

\*indicated that location was sampled for both 2012 and 2013, the other locations were sampled either in 2012 or in 2013.

To begin DNA extraction for sequencing, individual psyllid adults were placed into separate 1.5 ml microcentrifuge tubes with 150 µl tissue lysis buffer (10mM Tris pH=8; 50mM EDTA; 200mM NaCl; 1% (w/v) SDS), and ground for 1 minute using a pestle driven by a handheld electric mixer. Thereafter, DNA extraction was conducted following the instructions of the Qiagen DNeasy Blood & Tissue Kit (Qiagen, Valencia, CA). Aliquots of DNA from each insect were visualized on a 1.0 % agarose gel, with samples revealing a light band ~ 20 kp in size indicating that DNA was not degraded. The quantity of DNA extracted from each insect was measured using a Qubit 2.0 Fluorometer (Life Technologies, Grand Island, NY), which is highly selective for double-stranded DNA.

DNA samples were sent to SNPsauraus LLC (Eugene, OR) to generate NextRAD libraries and perform sequencing. To construct DNA libraries, genomic DNA (~10 ng) was first fragmented with the Nextera reagent (Illumina, San Diego, CA), which also ligated short adapter sequences to the ends of the fragments (Fig. S1, Supporting Information). DNA fragments were then amplified with two primers matching adaptor sequences, with one of the primers extending an additional nine nucleotides (GTGTAGAGC) as the selective sequence at the 3’ end (Fig. S1, Supporting Information). Thus, only fragments that could be hybridized to the selective sequence were efficiently amplified. Afterward, constructed DNA libraries were sequenced on an Illumina HiSeq2000 with 1x100 bp configuration to generate ~65X raw coverage.

### Sequence alignment, homology search and variant calling

The program Trimmomatic (Bolger *et al*. 2014) was used to remove the Nextera adapters and low quality reads (Phred quality score < 20). Thereafter, reads of all psyllids were pooled and aligned to each other to form an allelic cluster, and the read with the highest count in the population was chosen as a reference contig. In order to identify these contigs based on gene homology, we queried all 23,191 contigs to the NCBI reference sequence (RefSeq) database (Pruitt et al. 2007); we used the BLASTN algorithm from the BLAST program (Altschul et al. 1990) with an e-value cutoff of 0.0001. Subsequently, reads from each sample were aligned to the reference using the BWA-mem algorithm of the Burrows-Wheeler Aligner (Li & Durbin 2009). Variant sites were called using the mpileup and bcftools algorithms in SAMtools (Li et al. 2009). The missing genotypes were imputed, and genotype phasing was performed using the program Beagle 4.0 (Browning & Browning 2007).

### Inferring interrelatedness among potato psyllid subpopulations

The key goal of our study was to examine interrelatedness of potato psyllid subpopulations collected from bittersweet nightshade or potato sites, across a broad landscape, to infer possible movement patterns between the perennial and the annual host plant. We took four complementary approaches. First, we used a phylogenetic clustering technique in order to visually describe interspersion and/or separation of psyllids based on the host plant, site, and date from which the insects were collected. Second, we conducted an ancestry assignment analysis and principal component analysis to sort psyllids by likely ancestry and admixture, again allowing an examination of whether host plant, site and date of collection strongly aligned with ancestral groups and population structure. Third, as a means of inferring how patterns of psyllid interrelatedness differ through time and space, we calculated fixation indices (*F*_ST_) among pairs of collection sites and dates. Fourth, we utilized Analysis of Molecular Variance (AMOVA) to partition the genetic variance explained by host, time, ADMIXTURE clusters, and spatial separation within the main contiguous landscape we sampled (i.e., the spatial outgroup in Idaho was excluded). The details of each of these analyses are provided below.

#### Phylogenetic tree construction

We calculated genetic similarity (proportion of shared alleles) between all pairs of individuals using PLINK (v1.90; Purcell *et al*. 2007). We randomly selected one variant per contig so that the variants would be mostly independent. We then constructed a unrooted neighbor-joining tree (Saitou & Nei 1987) from the pairwise genetic distances using the nj function in the R package ape (v3.3; Paradis *et al*. 2004) and obtained support values by randomly resampling variants with replacement 500 times (bootstrapping).

#### Clustering analysis and population structure

We next used ADMIXTURE (Alexander *et al*. 2009) to delineate genetically-distinct groups within our psyllid collections, searching for the number of genetic lineages that best described the data. We increased the pre-defined number of ancestral populations (*K*) from *K* = 2 to *K* = 15. Ancestry coefficient matrices of *K* from 50 replicated runs were aligned and averaged using the program CLUMPAK (Kopelman *et al*. 2015). Because movement among sites could occur throughout our study region, and because ancestry and lineage of all psyllid populations was unknown, we conducted ADMIXTURE analysis in the “unsupervised” mode without providing any sampling information, and we identified the best *K* value as the run with the lowest cross-validation error (Alexander *et al*. 2009).

As a complementary approach examining the genetic population structure of psyllids separated by hosts, time and space, we also conducted principal component analysis (PCA) using the smartpca algorithm from EIGENSOFT (v6.0.1; Price *et al*. 2006).

#### F-statistics and spatiotemporal separation

We first estimated the inbreeding coefficient (*F*_IS_), based on Nei (1977), of psyllid collections that included ≥ 4 psyllids (Table S1, Supporting Information). Next we estimated population differentiation (*F*_ST_) between pairs of collections that included ≥ 4 psyllids (Table S1, Supporting Information), based on the equations described in Weir and Cockerham (1984) which take small and unequal samples sizes into account. First, we employed linear models to examine the relationship between geographic distance separating psyllid populations and their degree of genetic interrelatedness, working with insects collected between August, 2012 and October, 2013 (this was the time window during which the most sites were sampled roughly synchronously). Second, within sites sampled repeatedly through time, we examined the relationship between degree of temporal separation and the degree of genetic interrelatedness/divergence. All of the equations and calculations were implemented in R 3.1.2 (R Core Team 2013).

#### Analysis of Molecular Variance (AMOVA)

To compare the partitioning of genetic variance among hosts, and the partitioning among ADMIXTURE clusters, we ran two AMOVAs (Excoffier *et al*. 1992) on the Washington samples with collection population (location and time) nested within ADMIXTURE population or host plant (we used the poppr [Kamvar *et al*. 2014, 2015] and ade4 [Dray & Dufour 2007] R packages). Samples collected from the same site at different times were considered different collection populations. Samples from Washington were assigned to the genetic cluster with the greatest ancestry fraction as determined by the *K* = 3 run in ADMIXTURE. We tested significance for the host AMOVA using 1000 random permutations; significance was not assessed for the ADMIXTURE cluster run because testing the significance of clusters defined by exploratory analyses on the same dataset is circular (Meirmans 2015). To further explore the magnitude of genetic differences between groups of psyllids, we also performed pairwise *F*_ST_ analyses and calculated the median pairwise percent dissimilarity at all segregating sites.

## Results

### NextRAD sequencing, sequence alignment, homology search and variant calling

On average 400 Megabases of sequence data (in FASTQ format), equivalent to ~ 2.7 million 100 bp reads, were obtained from each psyllid NextRAD library. As no reference genome was available, we established a *de novo* assembly by combining cleaned reads across all samples. We searched for homologs of each contig in the assembly to the RefSeq database (Pruitt *et al*. 2007) in order to determine what possible genes or genomic regions these sequences might represent. However, only 2.7% (643 of 23,191) of the contigs returned homologies. Among the sequences that returned homologies, a large portion aligned with sequences of the Asian citrus psyllid, *Diaphorina citri*; *D*. *citri* is the most-closely-related insect species to the potato psyllid that has been sequenced (Reese *et al*. 2014). Because of such low percentages of likely homologous sequences, we did not pursue annotating the reference contigs. We identified 8,443 variants by aligning cleaned reads of each sample back to the reference.

### Inferring interrelatedness among potato psyllid subpopulations

#### Phylogenetic tree

Our neighbor-joining method demonstrated that samples taken from the geographically isolated bittersweet nightshade patch near Twin Falls, Idaho, formed a cluster separate from all other psyllids that we collected from either of the two host plants (Fig. 2). Otherwise, psyllids collected from potato fields in Washington were generally interspersed with insects collected from bittersweet nightshade patches in that same state (Fig. 2). An exception to this broader pattern was a group of psyllids collected from a suite of potato fields near Othello, Washington (Fig. 2, Table 1, Table S1, Supporting Information); this group of potato-collected psyllids fell out in a distinct clade separate from any psyllids collected from any other potato field, or from any bittersweet nightshade patch (Fig. 2). Note that the terminal branch lengths were quite long for this potato-collected “Othello” cluster, indicating substantial genetic variation among individual psyllids in the cluster (Fig. 2). Potato psyllids collected across two years at the Moses Lake bittersweet nightshade site clustered separately, suggesting genetic differentiation between years. In contrast, the insects from the two other bittersweet nightshade sites with multi-year collections (Mesa and Colfax) clustered together across the two years (Fig. 2).

**Fig. 2.**
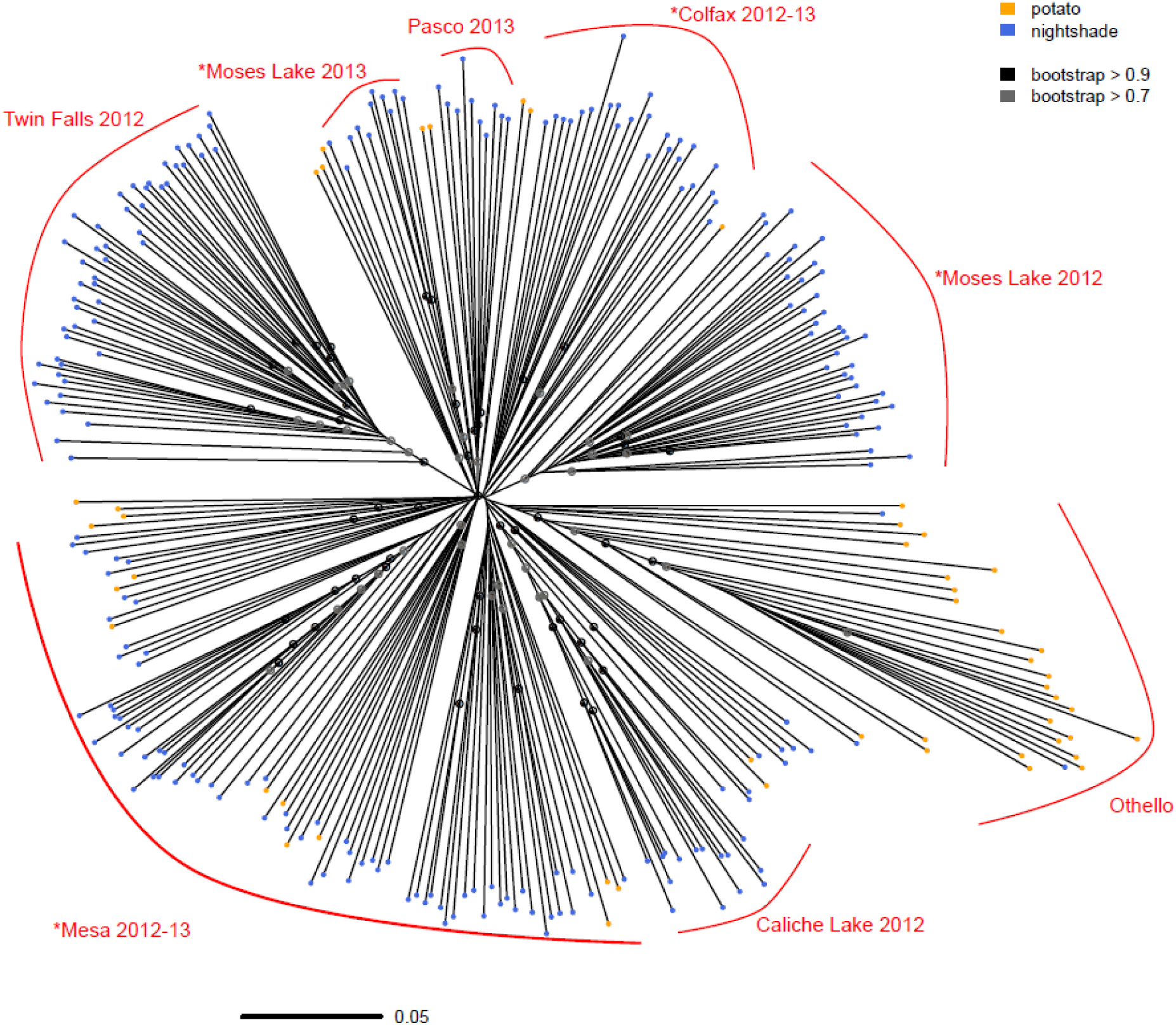
Unrooted neighbor-joining tree for potato psyllids collected from bittersweet nightshade patches (blue circles) or potato fields (yellow circles), constructed using proportion of shared alleles. Group labels were based on major clustering of the psyllids, do not include samples clustered into other clades due to admixture or gene flow. * indicates three nightshade locations with samples spanning two years.

#### Admixture and population structure

We represented clustering results of ADMIXTURE from *K* = 2 (minimum number of ancestral populations) to *K* = 8, where cross-validation error was lowest, suggesting the optimal modelling choice. At *K* = 2, we saw separation of the psyllids collected from our most geographically-distinct population, the single bittersweet nightshade patch sampled in southern Idaho, from all other psyllids collected on either of the two host plants (Fig. 3). The next group to separate from the others, at *K* = 3, was the same group of potato-collected psyllids, from potato fields near Othello, WA, identified in the phylogenetic tree as being genetically unique (Fig. 2, 3). *K* = 4 separated the potato psyllids collected from bittersweet nightshade at Caliche Lake; it also suggested genetic turnover for potato psyllids collected at the Moses Lake and Mesa sites between the two years during which those bittersweet nightshade patches were sampled, whereas the Colfax site exhibited constant genetic makeup across the two years (Fig. 3). *K* values of 5 through 8 identified relatively modest genetic divisions within sites and host-plant species.

**Fig. 3.**
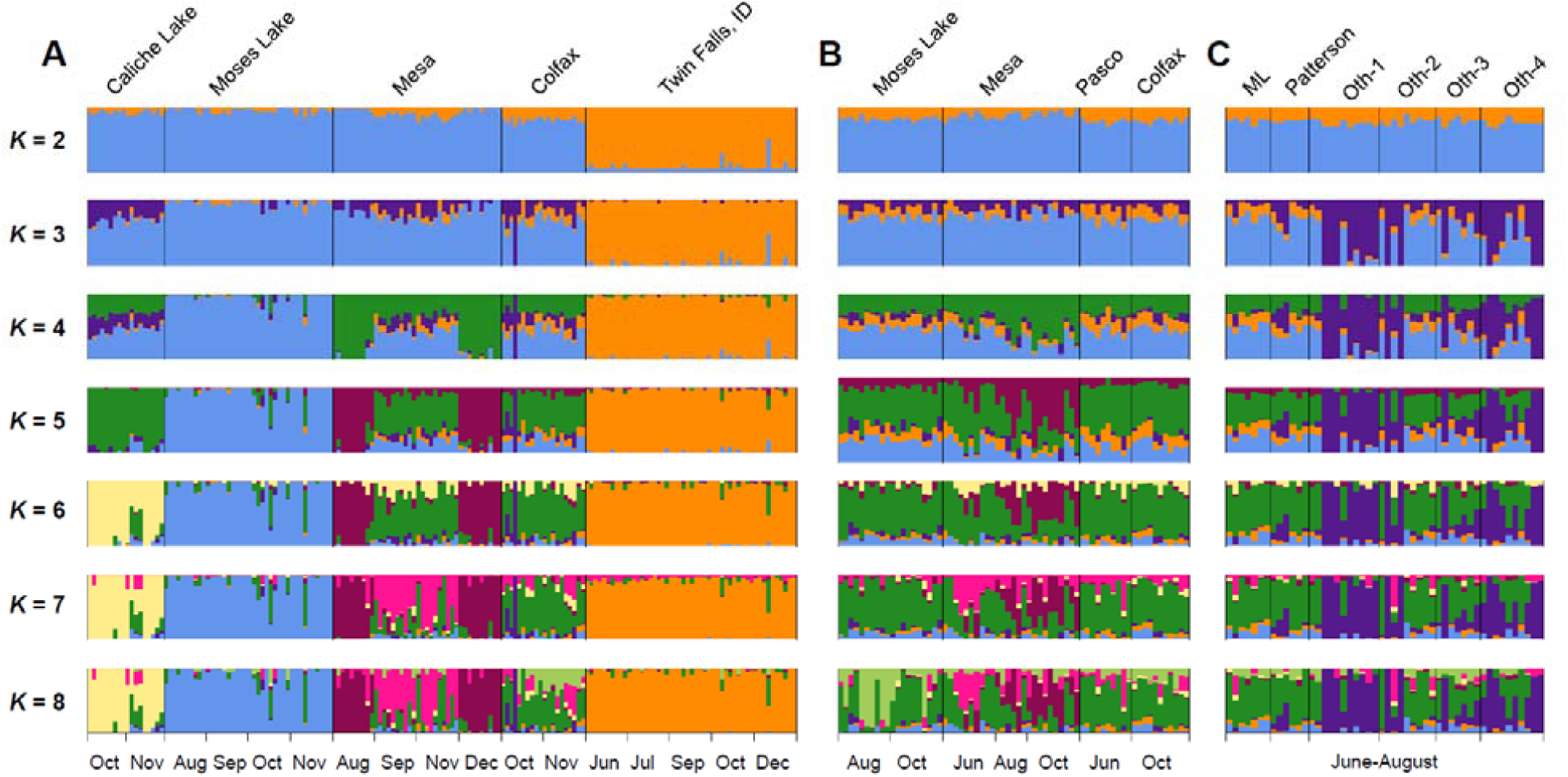
Estimated ancestry of potato psyllids collected from bittersweet nightshade patches in (A) 2012 or (B) 2013, or (C) from potato fields in 2013, with the number of ancestral populations (*K*) ranging from *K* = 2 to *K* = 8 (lowest cross-validation at *K* = 8, see Material and Methods). Ancestry coefficient matrices of *K* from 50 replicated runs were aligned and averaged using the program CLUMPAK (Kopelman *et al*. 2015). Each vertical bar represents a psyllid individual.

PCA revealed patterns similar to those detected with ADMIXTURE. Principal component (PC) 1 separated psyllids from Idaho from those collected in Washington, regardless of host plant (Fig. S2A, Supporting Information). Most potato-collected psyllids clustered with psyllids from nightshade sites in Washington, whereas some psyllids collected from potato fields near Othello, WA, clustered separately (Fig. S2B, Supporting Information). Patterns of temporal variation for psyllids from the three nightshade patches with multi-year samples reproduced what was found using ADMIXTURE (Fig. S2, Supporting Information).

#### F-statistics and geographic separation

The inbreeding coefficients (*F*_IS_) of psyllids collected from potatoes were markedly higher than the *F*_IS_ of psyllids from bittersweet nightshade patches (Wilcoxon signed-rank test, *p*-value = 0.0001, Fig. S3, Supporting Information), suggesting smaller effective psyllid populations in potato fields. We then investigated the correlation of population differentiation (*F*_ST_) and geographic distance. As described above, we included in these analyses the potato psyllids collected from August 2012 through October 2013, when insects were collected roughly synchronously across all sampled sites. We found a statistically-significant relationship between increasing degree of geographic separation and increasingly-large *F*_ST_ scores when the single Idaho bittersweet nightshade patch, the most-distant site that we considered, was included in the analysis (R^2^ = 0.43, df = 40, *p-*value = 9.6e-07; Fig. 3A). However, when that single-most-distant site was dropped from the analysis, this significant relationship disappeared (R^2^ < 0.1, df = 34, p-value = 0.8724; Fig. 3B).

We found a weak, but statistically-significant, correlation between within-site FST values and the duration of time between collections (R2 = 0.18, df = 48, p-value = 0.00199; Fig. 3C); this suggests increasing genetic differentiation within sites across time. In contrast to the relatively weak overall trend, FST of populations separated temporally at the Mesa and Moses Lake nightshade sites exhibited relatively large genetic changes through time. At the Mesa site within 2012, the FST of August and September populations was 0.134, compared to an FST between September and November of only 0.001 (Table S2, Supporting Information). At the Moses Lake site, the FST of any pairwise comparison between sampling dates within the same year was ≤ 0.021. In contrast, the FST of pairwise populations spanning two years was much higher (Table S3, Supporting Information), in agreement with the results of the phylogenetic tree and ADMIXTURE analyses. Psyllids at the Colfax nightshade site were more residential, as the FST of temporally-separated populations was constantly low (between 0.01 and 0.02 across the two years).

#### AMOVA

Among the many Washington sites, regardless of whether the top level was host or ADMIXTURE cluster, the largest component of genetic variability was explained at the individual level (Table 2), with little genetic differentiation among sites within genetic clusters and among individuals within sites. Consistent with the clustering analyses, relatively little genetic variation was explained by host plant, although the variation was significantly greater than zero (note that potato and nightshade populations were not always adjacent). However, a moderate amount of the genetic variation (~14%) was partitioned among ADMIXTURE populations (see Material and Methods). These results indicate that while overall there is little genetic differentiation between psyllids based on the host plant from which they were collected, the “third” genetic cluster identified by ADMIXTURE (purple bars in Fig. 2 associated with potato-collected Othello samples) is somewhat differentiated from the other Washington cluster. Nevertheless, the major source of variation is among individuals.

**Table 2.**
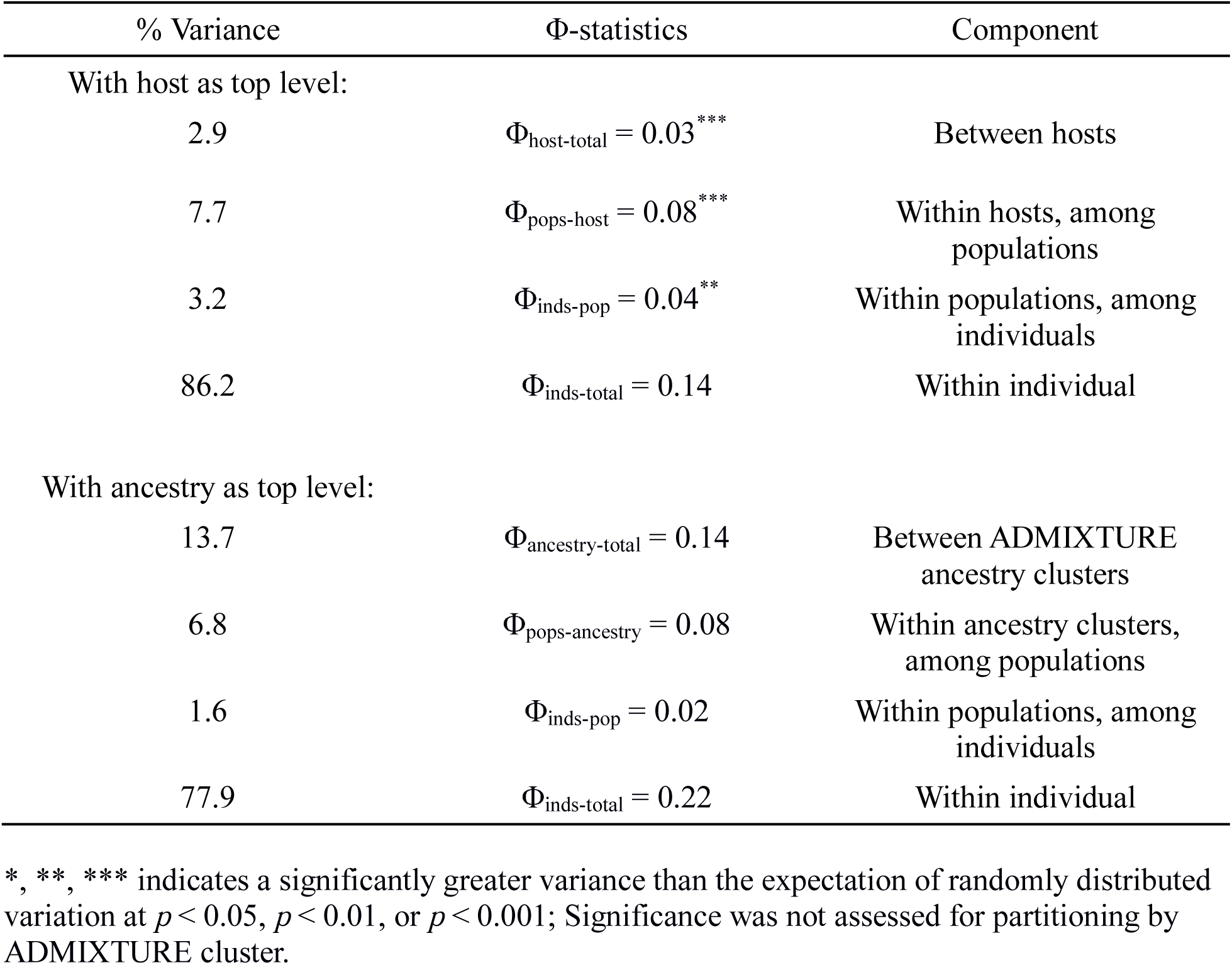
Results of two Analyses of Molecular Variance (AMOVAs) for samples within Washington. Populations were separated by sampling location and time. Individuals were assigned to the ADMIXTURE cluster with the greatest proportion of ancestry.

## Discussion

We took advantage of emerging NextRAD technology to examine interrelatedness of potato psyllids, vectors of a bacterium that causes zebra chip disease (Munyaneza 2012), collected from two host plant species. The insect has been suggested to overwinter on the perennial weed bittersweet nightshade before colonizing potato crops each summer, although this migratory linkage has never been demonstrated and many other putative non-crop hosts have been proposed (Horton *et al*. 2015a, Horton *et al*. 2015b). In most respects, we found that the psyllids from bittersweet nightshade and potato crops formed regularly interbreeding populations not clearly separated by host plant (Table 2, Fig. 2-3, Fig. S2, Supporting Information). For example, within our neighbor-joining tree (Fig. 2) potato-collected psyllids were generally interspersed among psyllids collected from bittersweet nightshade patches in the same region. Likewise, our ancestry analysis regularly showed potato- and nightshade-collected psyllids occurring with similar genetic profiles (Fig. 3). Finally, congruent PCA results showed a general interspersion of psyllids collected from the two host plants (Fig. S2, Supporting Information). All of these analyses are consistent with results from AMOVA, which assigned more variance to genetic differences between individuals than between sites or fields within a local region in Washington (Table 2). Overall, these results suggest that many psyllids are indeed moving from nightshade plants to potato crops as the latter are seasonally available. From an applied perspective, this suggests that removal of weedy, invasive bittersweet nightshade plants from the landscape might reduce a key source of potato psyllids eventually colonizing, and perhaps bringing the zebra chip pathogen to, potatoes.

At the same time, there was a second, genetically distinct group of potato psyllids found in four potato fields near Othello, Washington (Fig. 1, Table 1, Table S1, Supporting Information), that strongly differed from these overall patterns. Insects from those fields fell out as a unique clade in our neighbor-joining tree (Fig. 2), while clustering analysis suggested that this group was genetically distinct from the other psyllids on both potato and nightshade (purple bars from *K* = 3 through *K* = 8 in Fig. 3). A comparison of AMOVAs run with either host or genetic cluster as the top-level population indicated that about four times more of the genetic variation could be explained by genetic cluster than by host (Table 2). Furthermore, the magnitude of differentiation between the Othello group and the other Washington psyllids was greater than that between the other Washington psyllids and those from the distant Idaho site (Table S4, Supporting Information). There are at least two possible explanations for these findings. One is that there is a genetically-isolated sub-population of potato psyllids on bittersweet nightshade plants outside of our sampling network. An intriguing, second possibility is that a third host plant species is the source of the unique potato-collected insects, with insects on that as-yet-unidentified plant species genetically isolated from those on bittersweet nightshade. Many other putative potato psyllid host plant species have been suggested (e.g., other solanaceous weeds species, field bindweed *Convolvulus arvensis*, matrimony vine *Lycium barbarum*; Horton *et al*. 2015b). Thus it would be useful to collect potato psyllids from these other possible hosts and conduct the same NextRAD-based population genetics analyses reported here, to see if their genetic identity aligns with the distinct potato psyllids collected from the Othello potato fields. From an applied perspective, in turn, pest managers might consider the possibility that suppressing the exotic weed bittersweet nightshade might not entirely suppress regional potato-psyllid populations.

Several lines of evidence suggest that, despite the apparent stability of perennial bittersweet nightshade patches that may persist for decades, psyllids nonetheless regularly move between sites. For example, we observed genetic turnover between (and even within) years at our Moses Lake and Mesa sites, as evidenced by genetic differentiation seen in the ADMIXTURE analysis from *K* = 4 through *K* = 8 (Fig. 3). As a more general pattern, for pairings of collections within single sites but separated in time, we found that genetic divergence measured as *F*_ST_ values increased with increasing time between collection dates (Fig. 4C). This suggests a general, although relatively modest, turnover in genetic makeup across sites through time that would be consistent with movement among sites (although micro-evolutionary adaptation to particular sites could also explain this result; e.g., Watt *et al*. 2003). Furthermore, we noted no relationship between degree of genetic divergence and geographic distance between sites within Washington (Fig. 4B), consistent with frequent movement of insects among these sites. Geographic separation often strongly predicts genetic differentiation (e.g., Leebens-Mack & Pellmyr 2004; Massonnet & Weisser 2004), as indeed was the case when the most-distant Idaho site was included in analyses (Fig. 4A). Perhaps these potato psyllids move readily among sites in the absence of significant physical (e.g., the Blue and Bitterroot mountain ranges) and biological (e.g., the relative dearth of irrigated agriculture) barriers separating the Washington and Idaho sites. It remains unclear if the insects are moving for nutritional reasons (e.g., Belovsky 1981; Fraser *et al*. 1984; Wilmshurst *et al*. 1999), perhaps related to the seasonal drought typical of the region that could render irrigated potato crops more attractive than water-stressed bittersweet nightshade plants. Of course, a wide variety of other biotic (e.g., Long and Finke 2015) and abiotic (e.g., Canto *et al*. 2009) factors are known to trigger dispersal of pathogen-vectoring herbivores in other systems.

**Fig. 4.**
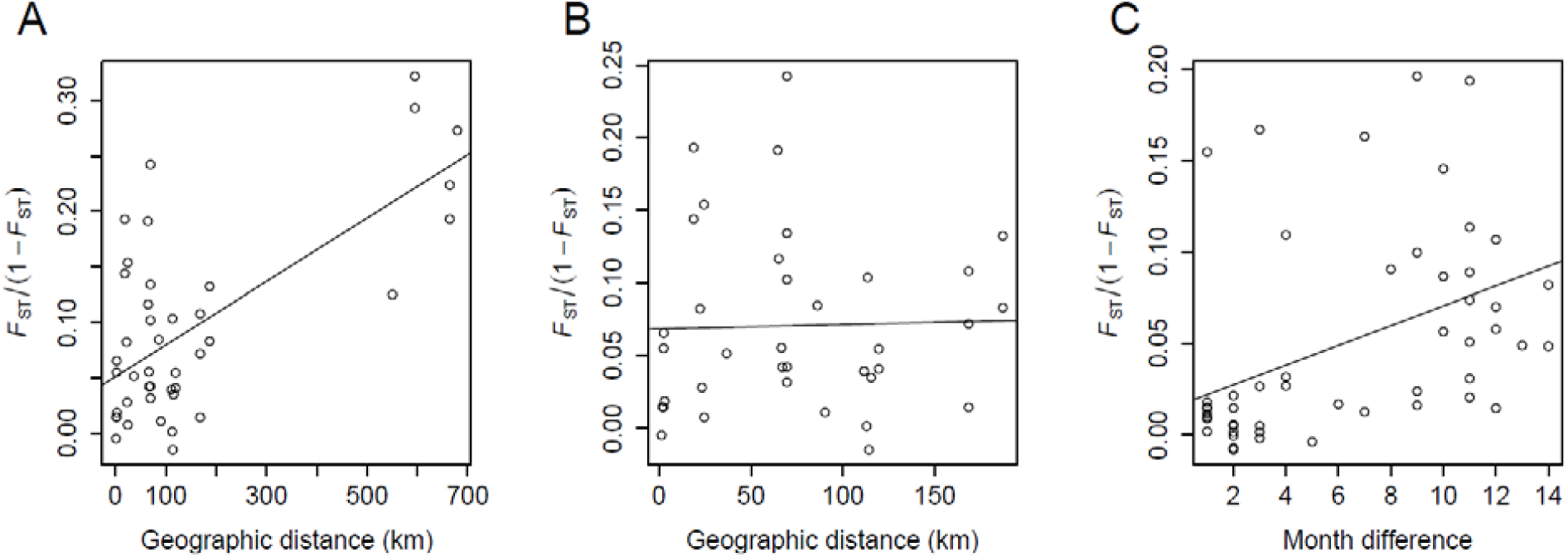
(A) Regression of *F*_ST_ values versus degree of geographic separation between pairs of psyllid populations differing in distance (but collected during the same month) for all population pairs across sampling dates from August, 2012 through October, 2013, and (B) for the same pairs of populations without the Twin Fall, ID, outgroup. (C) Regression of *F*_ST_ values for pairwise populations only differing in time (but collected at the same locations) across months.

While the findings we present here are specific to one particular plant-pathogen vector and a pair of its host plants, our approach could be widely applicable in other systems. Our study community is typical of many insect-vectored plant diseases, where a detailed understanding of local-scale movement is critical for understanding, predicting and managing plant disease dynamics. For example, outbreaks of bean leaf roll and pea enation mosaic viruses damaging to leguminous crops often depend upon movement of pea aphid (*Acyrthosiphon pisum*) vectors from alfalfa (*Medicago sativa* L.), a perennial host of both aphid and virus, onto peas (Clement 2006). This general movement of aphid and virus between these two host plants has been documented using microsatellite variation in the pea aphid (e.g., Eigenbrode *et al*. in revision ), a key and invaluable insight into understanding virus spread. However, sequencing approaches that detail SNPs across the genome, such as the NextRAD approach that we exploited, could delineate localized aphid and virus movement among particular fields within a growing region (e.g., Fig. 2, 3). In turn, this degree of resolution could provide field-specific predictions of disease risk that could be used by individual land-managers weighing treatment options. It is notable that fine-scale movement of vectors is important not just for predicting movement of plant pathogens, but also when highly-mobile vectors spread vertebrate pathogens. For example, the adult mosquitoes (*Anopheles* spp.) that vector devastating human pathogens such as Malaria (*Plasmodium* spp.) emerge from temporary water sources before traveling to reach human or other vertebrate hosts; for this reason understanding patterns of relatively short-distance movement between larval-development sites and human settlements is a critical part of accurately predicting disease outbreak (e.g., Thomas *et al*. 2013). Indeed, SNPs generated through RAD sequencing already hold promise for understanding relatively small-scale movement patterns of the mosquito *Aedes aegypti*, the vector of Dengue fever and other arboviruses (Rašić *et al*. 2014). This demonstrates the broad utility of these approaches for understanding the ecology of vector-transmitted diseases across diverse pathosystems.

The field of population genetics is increasingly making use of “Genotyping by Sequencing” to provide detailed genetic information on relatedness among populations. This approach has wide applicability in ecology and evolution, improving our understanding of site-specific adaptive evolution within species (Hohenlohe *et al*. 2010; Keller *et al*. 2013; Parchman *et al*. 2013) and evolutionary origins and dispersal patterns of migratory species (Larson *et al*. 2012; Hecht *et al*. 2013; Zhan *et al*. 2014), while helping to associate loci with particular phenotypic traits (Richards *et al*. 2013; Takahashi *et al*. 2013). While powerful (Hohenlohe *et al*. 2010; Hohenlohe *et al*. 2011; Richards *et al*. 2013; Lozier 2014; Rašić *et al*. 2014; Barley *et al*. 2015; Ebel *et al*. 2015; Szulkin *et al*. 2016), the most-commonly-used approach thus far, RAD sequencing, has limitations. Key among these is the reliance on restriction enzymes as a key part of the workflow, which limits the approach to use with vertebrates, or relatively large arthropods, from which a sufficiently-large quantity of DNA can be extracted from single individual animals (e.g., stickleback fish, Hohenlohe *et al*. 2010; land snails, Richards *et al*. 2013; butterflies, Ebel *et al*. 2015). NextRAD substitutes transposomes for restriction enzymes, necessitating less DNA per sample and thus allowing the approach to be used with smaller amounts of DNA than can be used with RAD sequencing (Russello *et al*. 2015). This is critical, because relatively small-bodied insects make up a majority of the most injurious herbivores of plants in many natural and agricultural settings (e.g., herbivorous mites, fruit flies, and whiteflies), while smaller arthropods serve as key vectors of some of the most damaging animal and plant pathogens (e.g., mosquitoes, ticks, fleas, aphids, and thrips). Using the NextRAD approach, we were able to describe relatively detailed patterns of interrelatedness among herbivorous insects that were too small to examine using earlier technologies. In turn, these patterns suggested local movement patterns of the vectors. We suggest that our work provides a model that may be of value in the many other systems where small-bodied insects move among host plants, and/or vector plant or animal pathogens.

## Acknowledgements

This research was supported by the National Institute of Food and Agriculture, U.S. Department of Agriculture, under Specialty Crop Research Initiative award number 2015-09273, along with grants from the Northwest Potato Research Consortium and the Washington State Commission on Pesticide Registration. C.I.C.C. was supported by a graduate research fellowship from Ecuador’s National Institute of Agriculture Research (INIAP) and the National Secretary of Higher Education, Science, Technology and Innovation (SENESCYT). We thank Carmen Blubaugh and Randall Svancara from Washington State University for their advice on the manuscript and assistance with data analyses on cluster computing, respectively. Finally, we thank Sanford Eigenbrode from the University of Idaho for his insightful suggestions on the manuscript.

## Data Accessibility

Raw DNA sequences were deposited into NCBI SRA with accession SRP062399.

## Author contributions

The study was designed by Z.F., C.I.C.C., A.S.J. and W.E.S., and manuscript writing was led by Z.F. and W.E.S. with assistance from B.E., J.L.K and A.O.B. Psyllid samples were collected by C.I.C.C., and DNA extraction was conducted by Z. F. Data analysis and interpretation were led by Z. F. and B.E with assistance from Q.Z. and A.O.B.

## References

Alexander DH, Novembre J, Lange K (2009) Fast model-based estimation of ancestry in unrelated individuals. Genome Research, 19, 1655–1664.

Altschul SF, Gish W, Miller W, Myers EW, Lipman DJ (1990) Basic local alignment search tool. Journal of Molecular Biology, 215, 403–410.

Bach CE (1980) Effects of plant density and diversity on the population dynamics of a specialist herbivore, the striped cucumber beetle, *Acalymma vittata* (Fab). Ecology, 61, 1515–1530.

Barley AJ, Monnahan PJ, Thomson RC, Grismer LL, Brown RM (2015) Sun skink landscape genomics: assessing the roles of micro-evolutionary processes in shaping genetic and phenotypic diversity across a heterogeneous and fragmented landscape. Molecular Ecology, 24, 1696–1712.

Behura SK (2006) Molecular marker systems in insects: current trends and future avenues. Molecular Ecology, 15, 3087–3113.

Belovsky GE (1981) Food plant selection by a generalist herbivore: the moose. Ecology, 62, 1020–1030.

Bischof R, Loe LE, Meisingset EL et al. (2012) A migratory northern ungulate in the pursuit of spring: jumping or surfing the green wave? The American Naturalist, 180, 407–424.

Bolger AM, Lohse M, Usadel B (2014) Trimmomatic: a flexible trimmer for Illumina sequence data. Bioinformatics, 30, 2114–2120.

Browning SR, Browning BL (2007) Rapid and accurate haplotype phasing and missing-data inference for whole-genome association studies by use of localized haplotype clustering. American Journal of Human Genetics, 81, 1084–1097.

Canto T, Aranda MA, Fereres A (2009) Climate change effects on physiology and population processes of hosts and vectors that influence the spread of hemipteran-borne plant viruses. Global Change Biology, 15, 1884–1894.

Chasen EM, Dietrich C, Backus EA, Cullen EM (2014) Potato leafhopper (Hemiptera: Cicadellidae) ecology and integrated pest management focused on alfalfa. Journal of Integrated Pest Management, 5, A1–A8.

Clement SL (2006) Pea Aphid Outbreaks and Virus Epidemics on Peas in the US Pacific Northwest: Histories, Mysteries, and Challenges. Online Plant Health Progress. doi:10.1094/PHP-2006-1018-01-RV

Coates BS, Sumerford DV, Hellmich RL, Lewis LC (2009) Repetitive genome elements in a European corn borer, *Ostrinia nubilalis*, bacterial artificial chromosome library were indicated by bacterial artificial chromosome end sequencing and development of sequence tag site markers: implications for lepidopteran genomic research. Genome, 52, 57–67.

Crowder DW, Northfield TD, Strand MR, Snyder WE (2010) Organic agriculture promotes evenness and natural pest control. Nature, 466, 109–112.

Despland E, Rosenberg J, Simpson SJ (2004) Landscape structure and locust swarming: a satellite’s eye view. Ecography, 27, 381–391.

Dray S, Dufour AB (2007) The ade4 package: implementing the duality diagram for ecologists. Journal of Statistical Software, 22, 1–20.

Ebel ER, DaCosta JM, Sorenson MD et al. (2015) Rapid diversification associated with ecological specialization in Neotropical Adelpha butterflies. Molecular Ecology, 24, 2392–2405.

Eigenbrode DS, Davis ST, Adams RJ et al. In revision. Not all herbivores are created equal: host-adapted aphid populations differ in their migratory patterns and capacity to colonize crops. Journal of Applied Ecology

Excoffier L, Smouse PE, Quattro JM (1992) Analysis of molecular variance inferred from metric distances among DNA haplotypes: application to human mitochondrial DNA restriction data. Genetics, 131, 479–491.

Fraser D, Chavez ER, Palohelmo JE (1984) Aquatic feeding by moose: selection of plant species and feeding areas in relation to plant chemical composition and characteristics of lakes. Canadian Journal of Zoology 62, 80–87.

Fryxell JM, Sinclair ARE (1988) Causes and consequences of migration by large herbivores. Trends in Ecology & Evolution, 3, 237–241.

Greenway G (2014) Economic impact of zebra chip control costs on grower returns in seven US states. American Journal of Potato Research, 6, 714–719.

Hagler JR, Jackson CG (2001) Methods for marking insects: Current techniques and future prospects. Annual Review of Entomology, 46, 511–543.

Hecht BC, Campbell NR, Holecek DE, Narum SR (2013) Genome-wide association reveals genetic basis for the propensity to migrate in wild populations of rainbow and steelhead trout. Molecular Ecology, 22, 3061–3076.

Hohenlohe PA, Bassham S, Etter PD et al. (2010) Population genomics of parallel adaptation in Threespine Stickleback using sequenced RAD tags. PLoS Genetics, 6, e1000862.

Hohenlohe PA, Amish SJ, Catchen JM, Allendorf FW, Luikart G (2011) Next-generation RAD sequencing identifies thousands of SNPs for assessing hybridization between rainbow and westslope cutthroat trout. Molecular Ecology Resources, 11, 117–122.

Horton DR, Cooper WR, Munyaneza JE et al. (2015a) A new problem and old questions: Potato psyllid in the Pacific Northwest. American Entomologist, 61, 234-244.

Horton DR, Cooper WR, Munyaneza JE et al. (2015b) Non-potato host plants of potato psyllid in the Pacific Northwest: a year-round complication? Potato Progress. Research & Extension for the Potato Industry of Idaho, Oregon, & Washington. XV, Number 2.

Illius AW, Clark DA, Hodgson J (1992) Discrimination and patch choice by sheep grazing grass-clover swards. Journal of Animal Ecology, 61, 183–194.

Kamvar ZN, Tabima JF, Grünwald NJ (2014) PopprU: an R package for genetic analysis of populations with clonal, partially clonal, and/or sexual reproduction. PeerJ, 2, e281.

Kamvar ZN, Brooks JC, Grünwald NJ (2015) Novel R tools for analysis of genome-wide population genetic data with emphasis on clonality. Plant Genetics and Genomics, 6, 208.

Keller I, Wagner CE, Greuter L et al. (2013) Population genomic signatures of divergent adaptation, gene flow and hybrid speciation in the rapid radiation of Lake Victoria cichlid fishes. Molecular Ecology, 22, 2848–2863.

Kopelman NM, Mayzel J, Jakobsson M, Rosenberg NA, Mayrose I (2015) Clumpak: a program for identifying clustering modes and packaging population structure inferences across K. Molecular Ecology Resources, 15, 1179–1191.

Koss AM, Jensen AS, Schreiber A, Pike KS, Snyder WE (2005) A comparison of predator and pest communities in Washington potato fields treated with broad-spectrum, selective or organic insecticides. Environmental Entomology, 34, 87–95.

Larson WA, Utter FM, Myers KW et al. (2012) Single-nucleotide polymorphisms reveal distribution and migration of Chinook salmon (*Oncorhynchus tshawytscha*) in the Bering Sea and North Pacific Ocean. Canadian Journal of Fisheries and Aquatic Sciences, 70, 128–141.

Leebens-Mack J, Pellmyr O (2004) Patterns of genetic structure among populations of an oligophagous pollinating yucca moth (*Tegeticula yuccasella*). The Journal of Heredity, 95, 127–135.

Li H, Durbin R (2009) Fast and accurate short read alignment with Burrows–Wheeler transform. Bioinformatics, 25, 1754–1760.

Li H, Handsaker B, Wysoker A et al. (2009) The sequence alignment/map format and SAMtools. Bioinformatics, 25, 2078–2079.

Liefting L, Sutherland P, Ward L, Paice K, Weir B, Clover G (2009) A New ‘*Candidatus* Liberibacter’ species associated with diseases of solanaceous crops. Plant Disease, 93, 208–214.

Long E Y, Finke DL (2015) Predators indirectly reduce the prevalence of an insect-vectored plant pathogen independent of predator diversity. Oecologia, 177, 1067–1074.

Loxdale HD, Lushai G (1998) Molecular markers in entomology. Bulletin of Entomological Research, 88, 577–600.

Loxdale HD, Lushai G (1999) Slaves of the environment: the movement of herbivorous insects in relation to their ecology and genotype. Philosophical Transactions of the Royal Society B: Biological Sciences, 354, 1479–1495.

Lozier JD (2014) Revisiting comparisons of genetic diversity in stable and declining species: assessing genome-wide polymorphism in North American bumble bees using RAD sequencing. Molecular Ecology, 23, 788–801.

Massonnet B, Weisser WW (2004) Patterns of genetic differentiation between populations of the specialized herbivore *Macrosiphoniella tanacetaria* (Homoptera, Aphididae). Heredity, 93, 577–584.

McElhany P, Real LA, Power AG (1995) Vector preference and disease dynamics: a study of barley yellow dwarf virus. Ecology, 76, 444–457.

Meglécz E, Anderson SJ, Bourguet D et al. (2007) Microsatellite flanking region similarities among different loci within insect species. Insect Molecular Biology, 16, 175–185.

Meirmans PG (2015) Seven common mistakes in population genetics and how to avoid them. Molecular Ecology, 24, 3223–3231.

Monteith KL, Bleich VC, Stephenson TR et al. (2011) Timing of seasonal migration in mule deer: effects of climate, plant phenology, and life-history characteristics. Ecosphere, 2, 1–34.

Munyaneza JE (2012) Zebra chip disease of potato: biology, epidemiology, and management. American Journal of Potato Research, 89, 329–350.

Nault LR (1997) Arthropod transmission of plant viruses: a new synthesis. Annals of the Entomological Society of America, 90, 521–541.

Nei M (1977) F-statistics and analysis of gene diversity in subdivided populations. Annals of Human Genetics, 41, 225–233.

Nextera DNA Library Prep Reference Guide https://support.illumina.com/downloads/nextera-dna-library-prep-reference-guide-15027987.html

Paradis E, Claude J, Strimmer K (2004) APE: Analyses of phylogenetics and evolution in R language. Bioinformatics, 20, 289–290.

Parchman TL, Gompert Z, Braun MJ et al. (2013) The genomic consequences of adaptive divergence and reproductive isolation between species of manakins. Molecular Ecology, 22, 3304–3317.

Peterson MA (1997) Host plant phenology and butterfly dispersal: causes and consequences of uphill movement. Ecology, 78, 167–180.

Power AG (1987) Plant community diversity, herbivore movement, and an insect-transmitted disease of maize. Ecology, 68, 1658–1669.

Price AL, Patterson NJ, Plenge RM et al. (2006) Principal components analysis corrects for stratification in genome-wide association studies. Nature Genetics, 38, 904–909.

Pruitt KD, Tatusova T, Maglott DR (2007) NCBI reference sequences (RefSeq): a curated non-redundant sequence database of genomes, transcripts and proteins. Nucleic Acids Research, 35, D61–D65.

Purcell S, Neale B, Todd-Brown K et al. (2007) PLINK: a tool set for whole-genome association and population-based linkage analyses. American Journal of Human Genetics, 81, 559–575.

R Core Team (2013) R: A Language and Environment for Statistical Computing. R Foundation for Statistical Computing, Vienna, Austria.

Rašić G, Filipović I, Weeks AR, Hoffmann AA (2014) Genome-wide SNPs lead to strong signals of geographic structure and relatedness patterns in the major arbovirus vector, Aedes aegypti. BMC Genomics, 15, 275.

Reese J, Christenson MK, Leng N et al. (2014) Characterization of the Asian Citrus Psyllid transcriptome. Journal of Genomics, 2, 54–58.

Redak RA, Purcell AH, Lopes JRS et al. (2004) The biology of xylem fluid-feeding insect vectors of *Xylella fastidiosa* and their relation to disease epidemiology. Annual Review of Entomology, 49, 243–270.

Richards PM, Liu MM, Lowe N et al. (2013) RAD-Seq derived markers flank the shell colour and banding loci of the *Cepaea nemoralis* supergene. Molecular Ecology, 22, 3077–3089.

Rose DJW (1979) The significance of low-density populations of the African armyworm *Spodoptera exempta* (Walk.). Philosophical Transactions of the Royal Society of London B: Biological Sciences, 287, 393–402.

Russello MA, Waterhouse MD, Etter PD, Johnson EA (2015) From promise to practice: pairing non-invasive sampling with genomics in conservation. PeerJ, 3, e1106.

Saitou N, Nei M (1987) The neighbor-joining method: a new method for reconstructing phylogenetic trees. Molecular Biology and Evolution, 4, 406–425.

Shariatinajafabadi M, Wang T, Skidmore AK et al. (2014) Migratory herbivorous waterfowl track satellite-derived Green Wave Index. PLoS ONE, 9, e108331.

Szulkin M, Gagnaire P-A, Bierne N, Charmantier A (2016) Population genomic footprints of fine-scale differentiation between habitats in Mediterranean blue tits. Molecular Ecology, 25, 542–558.

Takahashi T, Sota T, Hori M (2013) Genetic basis of male colour dimorphism in a Lake Tanganyika cichlid fish. Molecular Ecology, 22, 3049–3060.

Thomas CJ, Cross DE, Bøgh C (2013) Landscape movements of *Anopheles gambiae* malaria vector mosquitoes in rural Gambia. PloS One, 8, e68679.

Watt WB, Wheat CW, Meyer EH, Martin J-F (2003) Adaptation at specific loci. VII. Natural selection, dispersal and the diversity of molecular–functional variation patterns among butterfly species complexes (*Colias*: Lepidoptera, Pieridae). Molecular Ecology, 12, 1265–1275.

Weintraub PG, Beanland L (2006) Insect vectors of phytoplasmas. Annual Review of Entomology, 51, 91–111.

Weir BS, Cockerham CC (1984) Estimating *F*-Statistics for the analysis of population structure. Evolution, 38, 1358–1370.

Wilmshurst JF, Fryxell JM, Farm BP, Sinclair A, Henschel CP (1999) Spatial distribution of Serengeti wildebeest in relation to resources. Canadian Journal of Zoology, 77, 1223–1232.

Wink M (2006) Use of DNA markers to study bird migration. Journal of Ornithology, 147, 234–244.

Zhan S, Zhang W, Niitepõld K et al. (2014) The genetics of monarch butterfly migration and warning colouration. Nature, 514, 317–321.

Zhang D-X (2004) Lepidopteran microsatellite DNA: redundant but promising. Trends in Ecology & Evolution, 19, 507–509.

Zhu M, Radcliffe EB, Ragsdale DW, MacRae IV, Seeley MW (2006) Low-level jet streams associated with spring aphid migration and current season spread of potato viruses in the U.S. northern Great Plains. Agricultural and Forest Meteorology, 138, 192–202.

